# Single-tier point-of-care serodiagnosis of Lyme disease

**DOI:** 10.1101/2023.06.14.544508

**Authors:** Rajesh Ghosh, Hyou-Arm Joung, Artem Goncharov, Barath Palanisamy, Kevin Ngo, Katarina Pejcinovic, Nicole Krockenberger, Elizabeth J. Horn, Omai B. Garner, Ezdehar Ghazal, Andrew O’Kula, Paul M. Arnaboldi, Raymond J. Dattwyler, Aydogan Ozcan, Dino Di Carlo

## Abstract

Point-of-care (POC) serological testing provides actionable information for several difficult to diagnose illnesses, empowering distributed health systems. Accessible and adaptable diagnostic platforms that can assay the repertoire of antibodies formed against pathogens are essential to drive early detection and improve patient outcomes. Here, we report a POC serologic test for Lyme disease (LD), leveraging synthetic peptides tuned to be highly specific to the LD antibody repertoire across patients and compatible with a paper-based platform for rapid, reliable, and cost-effective diagnosis. A subset of antigenic epitopes conserved across *Borrelia burgdorferi* genospecies and targeted by IgG and IgM antibodies, were selected based on their seroreactivity to develop a multiplexed panel for a single-step measurement of combined IgM and IgG antibodies from LD patient sera. Multiple peptide epitopes, when combined synergistically using a machine learning-based diagnostic model, yielded a high sensitivity without any loss in specificity. We blindly tested the platform with samples from the U.S. Centers for Disease Control & Prevention (CDC) LD repository and achieved a sensitivity and specificity matching the lab-based two-tier results with a single POC test, correctly discriminating cross-reactive look-alike diseases. This computational LD diagnostic test can potentially replace the cumbersome two-tier testing paradigm, improving diagnosis and enabling earlier effective treatment of LD patients while also facilitating immune monitoring and surveillance of the disease in the community.

## Introduction

With the increased prevalence of emerging infections and vector-borne illnesses, it is critical to deploy robust and reliable testing platforms to combat the emergence and transmission of diseases^1^. Platforms that can be deployed rapidly and be used in point-of-care (POC) settings or for at-home testing can play a leading role in the rapid deployment of treatments for these diseases^2^. For example, during the COVID-19 pandemic, cost-effective rapid antigen tests and molecular diagnostic tests enabled quick isolation and therapeutic intervention for patients infected with the SARS-CoV-2 virus^3,4^. Lyme disease (LD) is a zoonotic infection caused by spirochetes of the *Borrelia burgdorferi sensu lato* complex that are transmitted through the bite of *Ixodes* ticks^5^. It is the most prevalent vector-borne disease in North America and Europe^6^ (Figure 1a). The incidence of the disease has continued to rise, exacerbated by climate change and the growing geographic distribution of tick populations^7^ (Figure 1b). LD is very difficult to diagnose using constitutional symptoms^8^ and there is no single definitive POC test currently available^9^. If not diagnosed and treated during early localized infection, the bacteria can disseminate to a variety of distal sites, resulting in serious tissue-type specific manifestations, including neurological, cardiac, or rheumatoid complications^10^.

**Figure 1.**
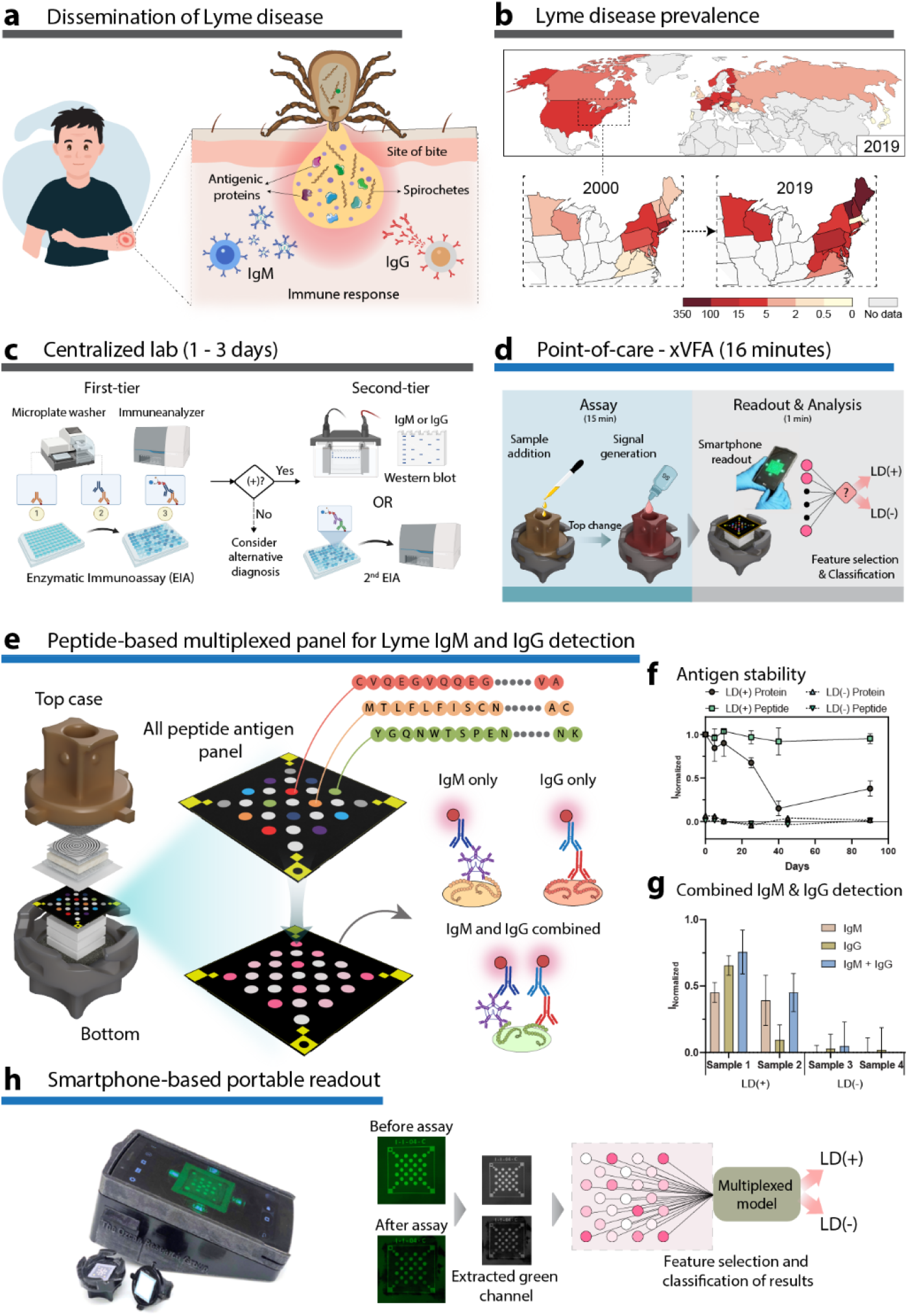
Overview of the paper-based multiplexed vertical flow assay (xVFA) and point-of-care diagnosis of Lyme disease (LD). **a** Transmission of Borrelia burgdorferi through the bite of Ixodeous ticks and the presentation of various antigens generating an immune response from the host. **b** Comparison of incidence of LD cases in the northeastern US from 2000 and 2019 indicating an increase in the incidence of cases due to the growing population of ticks. Worldwide incidence of LD in 2019. Legend indicates the number of cases per 100,000 people. **c** Centralized laboratory-based two-tier serology of LD uses relatively expensive instruments and trained personnel, resulting in high turnaround time and cost per test. **d** Point-of-care xVFA assay using low-cost paper layers and a smartphone reader that provides results for a multiplexed LD assay in <20 minutes. **e** The xVFA contains a selected peptide panel immobilized on a nitrocellulose membrane that reacts with IgM and IgG antibodies from LD patient serum. **f** Stability of modVlsE-FlaB peptide and VlsE recombinant protein indicating a loss in performance of a protein immobilized assay by more than 50% over a 90-day period. Standard deviation indicates three replicates. **g** Combining IgM and IgG detection in a single xVFA assay enhances the sensitivity of an individual immunoreaction spot. **h** Smartphone-based portable reader and automated image processing of the signals from the peptide panel before and after the assay, yielding normalized signal intensities. The individual peptide spots are analyzed using a multiplexed model to classify samples as either LD positive or negative.

Serological (antibody) testing of LD remains the mainstay approach for laboratory confirmation^11^. The U.S. Centers for Disease Control and Prevention (CDC) recommends a two-tiered approach for the diagnosis of LD, consisting of a first-tier enzyme immunoassay (EIA) which if positive or equivocal is followed by a second-tier western blot (WB) or a different EIA (Figure 1c). The two-tier system is widely understood to have significant drawbacks, requiring longer turnaround times^12^, underreporting of cases^13^, and a general failure to detect and treat LD in its early stages when treatment is most efficacious at preventing disseminated disease sequelae^14^. Furthermore, WB interpretation is subjective and the requirement of multiple specific protein bands to be positive results in failure in detecting most early-stage infections^15^. Additionally, commercial EIAs that are currently available use i) whole-cell lysates or recombinant proteins from single isolates of B31 species, which have cross-reactive epitopes that are common to other bacteria resulting in a high rate of false positivity^16^, or ii) single epitope-based detection, which precludes the recognition of antibodies to other immunodominant epitopes and does not take into consideration the variations in antibody production to different antigens over the time course of infection^17^. Further, these tests lack the level of multiplexing required to test for multiple antigens or the flexibility to include next-generation biomarkers clinically validated as potential diagnostic targets. Attempts to directly detect the pathogen using culture or molecular techniques have failed due to the transient presence of the bacteria in the bloodstream and low copy numbers of pathogen nucleic acids^18^.

Here we report a synthetic peptide based multiplexed vertical flow assay (xVFA), overcoming current limitations posed by traditional protein-based LD assays for point-of-care settings. The multiantigen panel consists of synthetic peptide-based immunogenic targets that detect total IgM and IgG antibody responses from patient serum samples in a single assay, resulting in a reduced cost and increased shelf-life. We used a machine learning-based diagnostic model to interpret the multiplexed results into a diagnostic recommendation enabling performance that matched the gold standard two-tier testing, all within a single POC testing platform. This peptide-based xVFA involves simple operational steps compatible with resource-limited settings such as rural tick-endemic regions. The multiplexed assay requires only 20 μL of serum sample and provides results in <20 minutes, limiting reagent and sample consumption and drastically improving the turnaround times for LD tests compared to currently available assays in widespread use. The work demonstrates the possibility of replacing traditional lab-based assays with robust multiplexed POC diagnostic platforms, promoting distributed healthcare systems and increased disease surveillance in the community. This is particularly relevant in the current public health landscape, with the COVID-19 pandemic highlighting the need for effective distributed diagnostic tools.

## Results

### Designing a single-tier POC assay for serologic testing of Lyme disease

The xVFA platform leverages a multiplexed array of immunoreactive peptide epitopes from *B. burgdorferi* and the ease of use of low-cost paper-based sensors to provide a single-tier, rapid and accessible platform to test for LD. Figure 1e-g illustrates an overview of the paper-based xVFA platform, which consists of multiple layers of paper with tuned flow properties that are stacked vertically. The different paper layers are assembled to ensure a uniform flow of samples and assay fluids across the entire cross-section of the sensing region, yielding an independent but relatively uniform environment for convection and reaction at each of an array of peptide spots. We were able to multiplex up to 25 immunoreaction spots with less than 8% CV and utilized a total of 9 different peptide antigens deposited in duplicates for all reported results unless otherwise stated (Figure 1e). We also found that using synthetic peptides has additional advantages of reduced cost and increased shelf-life when compared to full-length recombinant proteins of *B. burgdorferi* (Figure 1f). The modVlsE-FlaB peptide on the xVFA maintained > 95% reactivity over 90 days stored at room temperature, while full-length recombinant protein lost more than half of its reactivity under the same storage conditions and time.

To reduce the test complexity for POC use and increase its sensitivity for detecting both early and late LD patients, we evaluated whether combined IgM and IgG detection could be performed in the xVFA format. We screened a panel of secondary antibodies (Supplementary Table 1, Supplementary Figure S1) and found that by combining anti-IgM and IgG into a single assay, we could detect antibody binding to peptides at a higher rate than using IgM or IgG alone (Figure 1g). By combining the detection of IgM, which appears in higher concentrations in the bloodstream earlier post-infection along with IgG, the sensitivity of the LD test is improved. This is particularly important for early-stage disease diagnosis when current two-tier strategies have historically shown lower sensitivity.

Signals are read using a smartphone-based portable reader followed by automated processing of readouts to avoid bias in diagnostic interpretation (Figure 1h) ^19–21^. The custom reader consists of a tray that holds the paper assay device and slides into the reader, enabling the smartphone camera to capture images of the immunoreaction spots. Images are captured instantly and can be uploaded to the cloud for processing and automated analysis of signals. This feature can also promote interpretation of the results in a closed-loop setting, ensuring accurate reporting of tests to physicians and public health officials, and informing patient care in distributed health systems. The reader can be calibrated for use with any smartphone equipped with a camera and is able to capture images with optimal illumination using green LEDs. The use of green LEDs for illumination within the reader, combined with a controlled distance between the sensing layer and the smartphone camera lens, enables the device to consistently capture high-quality images regardless of user training.

### Epitope mapping and multiplexed panel design

Protein antigens contain a combination of linear and conformational epitopes that can be unique to a given organism (specific epitopes) or commonly found in other antigens present throughout the biosphere (cross-reactive epitopes). To limit the cross-reactivity of antigen targets, we performed linear B cell epitope mapping to identify epitopes specific to the *B. burgdorferi* proteins, OspC, DbpA, DbpB, BBK32, ErpP, p35, OppA2, RecA, LA-7, FlilB, BBA64, BBA65, BBA66, and BBA73. Figure 2a shows an overview of our investigation approach to identify unique antigen epitopes of *B. burgdorferi* that could be used as diagnostic targets in a multiplexed assay. Synthetic peptides containing proposed linear epitopes were screened by ELISA, either as a part of this study (Supplementary Figure S2) or in previous studies^16,22–26^, using panels of sera from Lyme disease patients, healthy volunteers, and patients with other look-a-like diseases (See methods, Sample cohort 1). In this study, we also evaluated peptides containing two epitopes from different antigens, modVlsE-FlaB, DbpA4-B6, and Var2FlaB which improved sensitivity in ELISA assays compared to individual peptides alone, while retaining the specificity by limiting non-specific binding^22, 24^. Peptides that demonstrated sensitive and or specific binding characteristics by ELISA were further screened using xVFA to determine if differences in the nitrocellulose membrane substrate or unique flow-through format affected antibody affinity.

**Figure 2.**
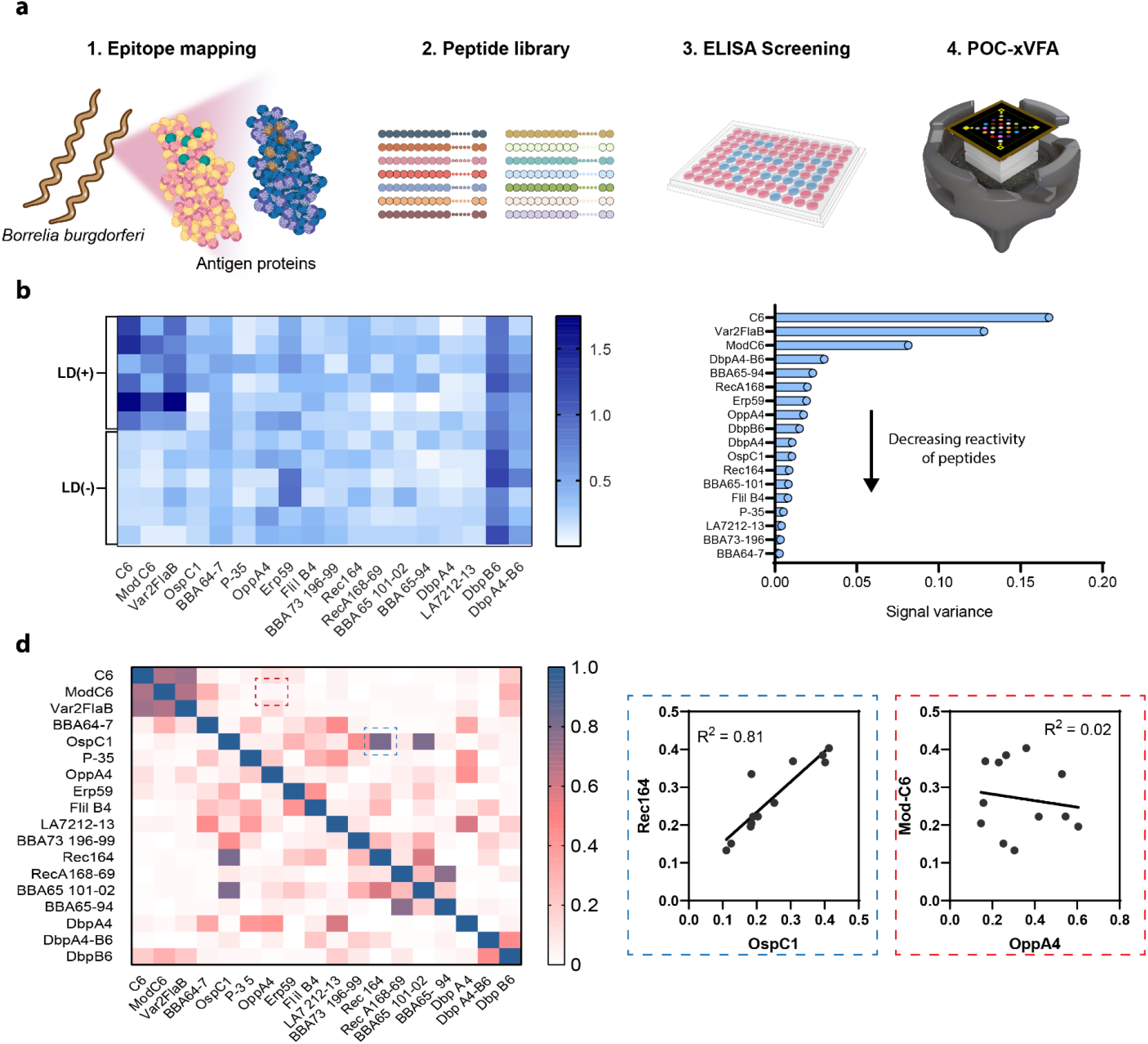
*B. burgdorferi* antigenic peptide library screening and selection of prevalent LD-specific epitopes. **a** Overview of the screening of epitopes to select the most relevant peptide antigens that function in the paper-based xVFA platform. **b** Heatmap representing the reactivity of peptides screened against LD positive (+) and negative (-) patient samples to develop the paper-based multi-antigen xVFA platform. **c** Variance of the peptide signal intensity obtained using xVFA plotted in decreasing order of reactivity to identify the most active antigens. **d** Heatmap representing the correlation of peptides against each other. Inset shows examples of highly correlated peptides (Rec164 and OspC1) and non-correlated peptides (modVlsE-FlaB and OppA4) which yield additional information for a diagnostic panel.

We measured the reactivity of the peptides immobilized on the nitrocellulose sensing membrane using a panel of well-characterized serum samples containing six LD-positive samples from different stages of the disease and six healthy control samples (See methods, Sample cohort 2). Figure 2c shows a heatmap of the signals corresponding to the signal detected for each peptide spot. Peptides modVlsE-FlaB and Var2FlaB showed 100% sensitivity and specificity. DbpB6 and DbpA4-B6 which is a dipeptide combining epitopes from both DbpA4 and Dbp-B6, showed high reactivity even in healthy control samples, indicating non-specific interactions that were not observed in the ELISA format. Peptides Erp59, OppA4 and BBA64-7 showed reactivity with four out of six positive serums tested, while also demonstrating non-specific interactions with one out of the six non-LD sera. A list of all peptides that were screened using the xVFA can be found in Supplementary Table 2. Figure 2D shows the activity of the peptides as measured by the variance in the signal across both the LD positive and negative samples. As previously identified, DbpA4-B6 and DbpB6 spots had high variance in signal across patients, but this high variance was also connected to high background interactions against healthy samples and therefore was not selected for further test development. By plotting the correlation of the peptide signals across patients (Figure 2E) we identified and removed correlated epitopes where antibodies against both epitopes were high or low in tandem and were already represented on the multiplexed panel. For example, both OspC1 and Rec164 were moderately reactive across LD patients and had a higher degree of correlation (R^2^=0.81). Therefore, only OspC1 was included in the multiplexed panel as it is expressed early during the transmission of disease and is a marker for early-stage infection. Further, OppA4 was given higher importance over other peptides as it was reactive in LD patient sera and did not correlate with binding to other candidates (R^2^< 0.4 to all other peptides).

Following the initial analysis of antibody binding to 18 peptides, we down-selected to a subset of 9 peptides, which when spotted in duplicate along with 3 positive and 4 negative internal controls, would provide for a total of 25 immunoreaction spots on our array. As discussed, this subset was selected based on antibody binding signal levels in LD patients vs. controls and was prioritized for those that were not correlated to signatures for other peptides in the pool. This expanded positive readouts to a larger fraction of patients to increase sensitivity. Thus, it could be used to capture additional epitope-specific information that was not obtained from other peptides. Based on the criteria of reactivity and uniqueness, we selected 9 peptides: modVlsE-FlaB, Var2FlaB, BBA64-7, OspC1, OppA4, p-35 21, ErpP59, Flil-B4, and BBA73 196-99 for the training of our final xVFA diagnostic model with clinical samples.

### Training of xVFA peptide panel using early-stage LD samples

To train the deep-learning diagnostic model, a cohort of 50 serum samples (See methods, Sample cohort 3) was used. The cohort included 25 early-stage LD samples and 25 healthy control samples collected by the Lyme disease biobank (LDB) from LD endemic regions (Supplementary Table 3). Nine peptide targets from 10 different antigens of *B. burgdorferi* were used to start the training of the diagnostic model. Peptides were spotted in duplicate with positive and negative controls (Figure 3A). The mean signal intensity averaged over the two spots from three separate assay replicates shows distinct peptide reactivity signatures that vary between LD and healthy control samples in the training cohort (Figure 3B). Notably, the test showed reproducibility - the coefficient of variation (CV) of the average spot intensity per peptide across the three replicates was always below 10% (Figure 3B).

**Figure 3.**
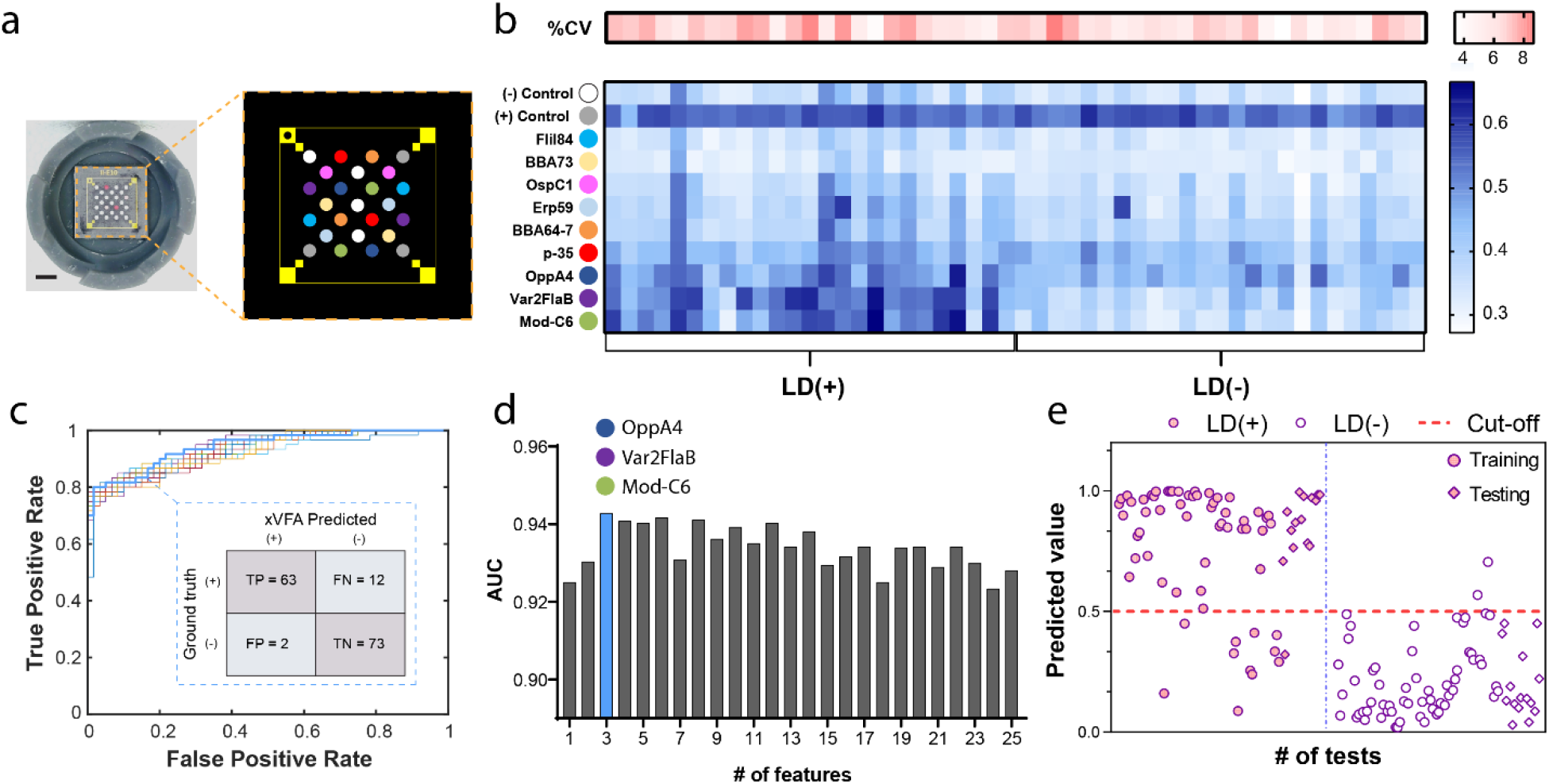
Training of the multi-antigen xVFA panel using early-stage LD patient samples obtained from the Lyme Disease Biobank. **a** The multi-antigen coated sensing membrane and map of each antigen location spotted in duplicates along with the positive and negative control reaction spots. **b** Heatmap representing the average signal intensity obtained from the multi-antigen panel for the individual peptides and the positive and negative control spots. The color scale on the top represents the average %CV in measurement for all peptides per patient sample. **c** Receiver operator characteristics (ROC) resulting from the neural network-based multiplexed diagnostic model comparing the model’s performance when different numbers of peptides are used to train the network. The inset shows the confusion matrix for a 3-peptide model. **d** Bar plot showing the area under the curve (AUC) for the different input features used to train the diagnostic model. A 3-peptide model including single immunoreaction spots from modVlsE-FlaB, Var2FlaB, and OppA4 yields the highest AUC of 0.94. **e** The final prediction outcome from the neural network-based diagnostic model, indicated as values from 0 to 1, where each dot represents a test that was perform on the xVFA using a patient sample (50 patient samples tested in triplicate). The red dotted line at 0.5 indicates the threshold for positivity of the diagnostic model.

We first investigated whether the signal from a single peptide would provide sufficient diagnostic performance for LD testing. The use of individual peptides in isolation fails to meet the performance requirements to serve as a single-tier test for Lyme disease diagnosis, with the best performance being 76% sensitivity and 95% specificity for the modVlsE-FlaB peptide (Supplementary Figure S3). We then turned to a deep learning-based analysis of the multiplexed signals to form a synergistic model that combined information from the spots into a binary classification of either positive or negative detection.

A diagnostic model was trained using a deep-learning neural network to classify a sample as LD positive (prediction value > 0.5) or negative (prediction value ≤ 0.5) using a cross-validation approach where 120 tests (40 patients × 3 replicates per patient) were used to optimize the architecture of the neural network model (see Methods) and 30 tests (10 patients × 3 replicates per patient) were used to validate the optimized model. We used the model to first select the next most important immunoreaction spot to improve diagnostic accuracy as defined by the area under the curve (AUC) of the receiver operating characteristic (ROC). Figure 3C shows the peptide selection process using sequential forward feature selection (SFFS) and a comparison between the performance of different combinations of peptides as shown by the ROC curves. Here, each immunoreaction spot is considered a feature that serves as an input to the diagnostic model, totaling 25 different immunoreaction spots for nine peptide antigens plus control spots. The AUC for the different feature combinations is shown in Figure 3D and yields a maximum when using a combination of three peptide immunoreaction spots: modVlsE-FlaB, Var2FlaB and OppA4. Using the combination of these three peptides, we achieved a sensitivity of 84% and a specificity of 97.3% which exceeded the performance of the individual peptides or antigens in a standalone assay. modVlsE-FlaB, Var2FlaB and OppA4 each yielded a sensitivity of 76%, 76% and 7%, respectively with specificity set at 95% for each. Figure 3E shows the prediction of our multiplexed diagnostic model using the three peptides selected with the SFFS method. Each dot represents the model output value for a separate xVFA test, where a threshold for positivity is set at 0.5. Samples used for training and testing are separately indicated on the graph. A clear separation is observed in model outputs for LD positive and negative patients for both training and testing sets. Of the 3 samples (LD50, LD72, LD74) with a false negative prediction (<0.5) gold standard testing also confirmed a weak antibody response. These three LD+ patient samples were from patients who did not present with an EM at the time of enrollment and remained negative or indeterminate by the IgG western blot. Additionally, two other positive samples (LD31, LD61) were negative for the C-6 peptide ELISA and for the IgG western blot but was found positive using xVFA. This suggests that having multiple epitopes represented can improve upon sensitivity when traditional single target tests are not effective. Two replicates from two different healthy control samples were misclassified as positive with a prediction value that was closer to the 0.5 threshold. The samples had higher background interaction with all the peptides resulting in a final prediction value that was higher than the threshold of 0.5. The xVFA was able to correctly identify the samples from the training cohort as either LD positive or negative with higher accuracy (94%) compared to the first-tier centralized lab based C6 peptide ELISA (Oxford Immunotec, Marlborough, MA) with an accuracy of 92% among the same samples tested. Additionally, the xVFA also outperformed the second-tier IgM and IgG western blot (Viramed; Biotech AG, Germany) which had an accuracy of 84% and 66% respectively.

### Clinical assessment by blinded testing

Using the optimal diagnostic model outlined earlier, we evaluated the performance of our single-tier multiplexed POC test using serum samples obtained from the CDC Lyme disease repository in a blinded fashion in triplicate (Figure 4a). The cohort consisted of a diverse set of samples collected from patients with different stages of LD, healthy control groups from both endemic and non-endemic regions, and patients with look-alike disease and no previous exposure to LD. Overall, our POC xVFA performed similarly (no statistically significant difference) compared to the lab-based standard two-tier diagnostic testing for LD but had the advantage of being single-tier and being performed in a rapid POC format that could be interpreted during clinical evaluation of patients (Figure 4b). All four of the samples corresponding to acute-stage infections were classified as LD negative using the standard two-tier serology algorithm (EIA and Western Blot) and the xVFA diagnostic model. However, the diagnostic model achieved 100% specificity across all categories of healthy control groups, including zero false positives for look-alike diseases. High specificity in distinguishing look-alike diseases and healthy control samples distinguishes our xVFA approach from other single biomarker-based commercial platforms that showed cross-reactivity with Syphilis, an infection caused by a spirochete, and also exhibited cross-reactivity with healthy control samples from both endemic and non-endemic regions (Supplementary Table 4). By selecting specific *B. burgdorferi* epitopes, training a model to identify the most useful epitopes in the context of xVFA, and using early-stage LD samples to refine the model we were able to successfully expand detection sensitivity while eliminating cross-reactive epitopes and improving the diagnostic outcome towards a single-tier multiplexed POC assay.

**Figure 4.**
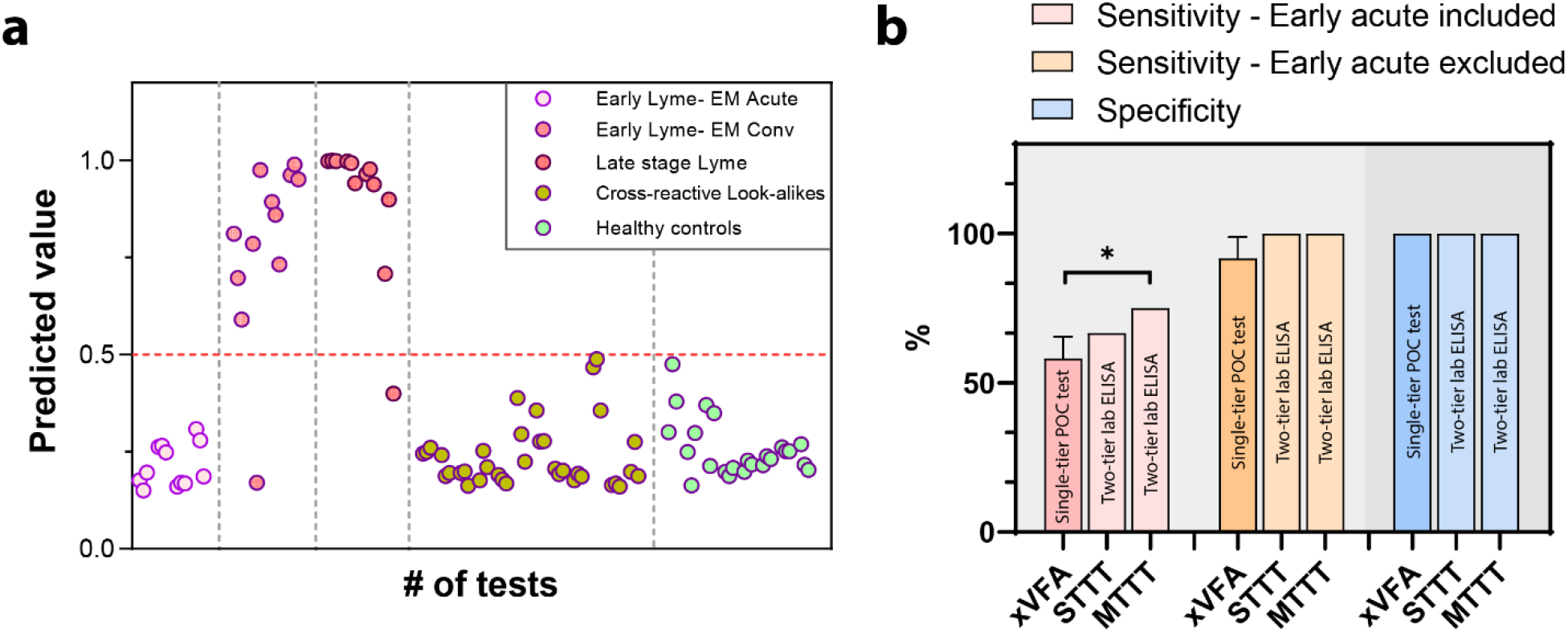
Blinded validation of the multi-antigen xVFA and the neural network-based diagnostic model using patient samples obtained from the CDC. **a** Prediction outcome from the neural network-based diagnostic model using the trained network indicated as values from 0 to 1, where each dot represents a test that was performed on the xVFA using patient samples that had different stages of LD, healthy control samples from regions where LD is endemic and non-endemic, cross-reactive samples from look-alike diseases such as fibromyalgia, rheumatoid arthritis, multiple sclerosis, mononucleosis, syphilis, and severe periodontitis. The red dotted line at 0.5 indicates the threshold for positivity of the diagnostic model. Three xVFA tests were run per sample. **b** Bar plot comparing the sensitivity (with and without the inclusion of early acute stage samples) and specificity of the rapid point-of-care, multiplexed vertical flow assay (xVFA) with centralized lab based Standard Two-tier Testing (STTT) and Modified Two-tier Testing (MTTT), gold standard diagnostic algorithms commercially used for LD diagnosis. One sample t-test was performed to compare the performance of the xVFA and the two-tier gold standard algorithms. The xVFA sensitivity and specificity was not statistically significantly different (*p>0*.*05*) from the two-tiered centralized lab assays with the exception of MTTT algorithm (*, *p<0*.*05*) when early-stage disease samples were also included.

## Discussion

Synthetic peptides are excellent capture antigens as they preserve the binding sites for the detection of anti-*B. burgdorferi* antibodies while reducing the cost and complexity of diagnostic tests. Peptides can be chemically synthesized at extremely low-cost and on a scale to manufacture tests in bulk. Native, or recombinant proteins of *B. burgdorferi* are harder to synthesize, purify and isolate, leading to higher costs, particularly in downstream quality control (Supplementary Table 5), and have a reduced shelf-life before test performance is impacted (Figure 1f). Recombinant proteins may also contain cross-reactive epitopes, which limit the performance, reducing specificity, as seen in our previous study using a multiplexed panel of *B. burgdorferi* antigens and a single peptide^27,28^. Our POC xVFA using the three peptide antigens identified by our model is projected to cost significantly less than traditional EIAs while achieving superior diagnostic performance. The material cost associated with a single xVFA is less than $3 and can be manufactured at scale owing to compatibility with automated spotting equipment commonly used in LFA production (Supplementary Table 5). This would provide greater access to testing in resource-limited and rural areas.

By selecting epitopes that are highly conserved among different strains of *B. burgdorferi* with limited homology to sequences in other antigens likely to be encountered by humans we can reduce cross-reactivity. Through the inclusion of multiple epitopes with our machine learning framework, we can maintain or improve sensitivity compared to current assays, while having a limited epitope pool compared to a whole recombinant protein or bacterial sonicates. Starting with a set of 18 curated peptides, we further down-selected using our initial screening data set to eliminate peptides that did not bind well to paper, were highly correlated to other peptides, or were generally reactive to LD negative sera. With a smaller set of 9 peptides, we screened a cohort of 50 samples to develop a training data set for further refinement of a diagnostic model to maximize sensitivity while avoiding false positive results. Our final model selected 3 immunoreaction spots (3 different peptides, modVlsE-FlaB, Var2FlaB and OppA4, representing 4 different antigens).

Our POC xVFA had no significant difference in performance compared to the centralized lab based two-tier approach. When used as a single-tier assay the xVFA had 100%, 87.5% and 87.5% agreement across the three replicates with both standard and modified two-tier testing results from the CDC Lyme disease repository, a sample set the model had never seen (Figure 4b). Although current lab-based EIAs are used as part of a two-tier assay, if EIAs are used as single-tier tests there was a tradeoff of sensitivity vs. specificity in comparison to the xVFA. For example, commercial ELISA tests such as Zeus IgG did not detect any of the acute samples (or convalescent samples) but were very specific with no cross-reactive binding. On the other hand, the Zeus IgM EIA, WCS EIA and other Zeus VlsE/pepC10 EIA all detected one or two of the acute-stage samples but were also found to non-specifically bind to look-alike disease samples and healthy patient samples from endemic and non-endemic areas. The high specificity of our xVFA combined with the enhanced sensitivity in a POC format would be suitable in a rapid, POC test setting such as local pharmacies, clinics, or other POC settings, enabling healthcare providers to quickly determine appropriate treatment with antimicrobials upon obtaining a positive result. A negative result may indicate repeat testing is needed in acute patients with other signs and symptoms after waiting for serum IgM and/or IgG antibody levels to increase further. The engineering of xVFA and the underlying immunochemistry results in a highly repeatable test with a CV of 8.5% between different tests by the same operator and a CV of 9.3% for tests performed by different operators who are not specialized in clinical testing (undergraduate students), ensuring repeatable results independent of operator training.

The utilization of a selected subset of three peptides in the xVFA implies that the majority of immunoreaction spots on the platform are not utilized. Future work could investigate whether replacing these spots with copies of the selected peptides would lead to even further improved performance. Alternatively, epitopes for other tick-borne illnesses or other related bacterial pathogens could be included to expand the diagnostic capabilities or monitor other epitope-specific antibodies from patients and acquire additional data on the prevalence of different immunophenotypes across a significantly expanded set of patients. Using diagnostic tests to derive epitope-specific community health information would serve to understand the underlying disease yielding critical information on LD pathogenesis and better formulation of treatment measures. Large data sets that leverage multi-epitope arrays associated with clinical outcomes could also be used to train even better models to diagnose LD or develop region-specific models, if required.

Cloud connectivity with a smartphone implementation can enable the integration of Lyme diagnostic results with patient care and public health guidance. The monitoring of new positive disease incidences could also enable tracking of tick-borne diseases informing public health guidance in areas of high endemicity. Smartphone-based interpretation also eliminates bias in results interpretation by serving as a seamless interface between the diagnostic test and the neural network-based prediction model. The reported test combined with recent advances in telemedicine and at-home diagnostics, coupled with smartphone-based interfaces, could enable more efficient test-to-treat paradigms, seamlessly integrating physician input for prescription, therapeutic delivery, and ultimately leading to more rapid and effective patient care.

## Methods

### Overview of the multiplexed vertical flow device (xVFA) platform

The vertical flow assay platform consists of paper layers that are stacked together to form a 3-dimensional fluidic network that transports assay fluids vertically by capillary wicking action. The paper layers are housed inside a plastic cassette that can be separated into two parts through a twist mechanism, revealing a multiplexed sensing membrane on the top of the bottom section, containing 25 immunoreaction spots spatially isolated by a hydrophobic wax-printed barrier. Each spot is preloaded with different capture peptides (1 mg/mL) and a Goat anti-Mouse IgG (0.1 mg/mL) that acts as the positive control spot for the assay. The top section of the device contains paper layers that are engineered to ensure uniform flow of samples and reagents across the entire cross-section of the sensing membrane. The bottom consists of highly efficient absorption pads below the sensing membrane that act as sink for the assay fluid. For each assay, we use two top sections of the device during operation. The first top introduces the sample into the sensing membrane followed by incubation, which ensures binding of *B. burgdorferi* specific antibodies to the different peptide antigens immobilized. The second top is used for signal generation by addition of anti-human IgM and IgG antibodies conjugated to gold nanoparticles (AuNPs). The AuNPs bind to the LD specific IgM and IgG antibodies that were captured and the non-specific signals are washed away by buffer solution. The xVFA operation was optimized by tuning the running buffer components, sample volume, and pore size of the nitrocellulose membrane using control LD positive and healthy samples (Supplementary Figure 4). Upon completion of the assay, the sensing membrane is imaged using a low-cost portable smartphone reader that captures the individual reaction intensities under a green illumination for maximum light absorbance and signal-to-noise. The raw intensities from the reaction spots are then normalized and analyzed using an image processing algorithm and multiplexed diagnostic models to determine a final verdict on seropositivity. The entire assay and the diagnostic inference based on the different immunoreaction spots can be completed in under 20 minutes. The overall material cost for the entire consumables is less than $0.4 per test (Supplementary Table 5).

### Assay operation

To perform the xVFA assay, the first step is to record a background image of the unused sensor using the portable smartphone reader. The device is then assembled by attaching the first top case to the bottom case with a simple twist and activated by adding buffer solution and 20 μL of serum sample. After the buffer and serum are fully absorbed into the xVFA cassette (8 minutes), the first top case is opened and replaced with the second top case for signal generation. The 40 nm AuNPs-conjugated detector antibody solution (a mixture of mouse anti-human IgM and IgG in a 1:1 ratio) and running buffer are introduced into the device and incubated for an additional 8 minutes. During this time, the AuNP conjugated to the detector antibodies will react with human anti-borrelia immunoglobulins, producing an intense signal. Once the incubation period is complete, the xVFA cassette is opened and an image of the multiplexed signal on the paper membrane is captured using the mobile-phone reader.

### Smartphone reader

The portable assay reader consists of a smartphone (LG G4H810) with a 3D printed opto-mechanical attachment that contains four 525 nm wavelength light-emitting diodes (LEDs) for uniform illumination of the sensing membrane. An external lens is mounted below the built-in camera lens of the smartphone within the 3D printed attachment. All images were taken in raw dng format using the standard Android camera app on the smartphone. To measure the signals, the bottom case is connected to a 3D printed tray and slid into the reader for easy and repeatable measurements. The cost of the reader components combined with the smartphone is less than $200.

### Image processing and neural network-based analysis

To analyze the results of the assay, raw dng images of the sensing membrane taken before (background image) and after (signal image) the assay is first converted to tiff format. The green pixels are then extracted, and the background and signal images are registered to each other using a rigid transformation. The immunoreaction spots are identified in the background image, and a fixed-radius mask is defined for each spot, covering approximately 80% of the immunoreaction spot area. The pixel intensity within this mask is then calculated for the registered signal image and normalized by the average pixel intensity of the corresponding immunoreaction spot in the registered background image. This background normalization procedure helps to compensate for nonuniformities in illumination and local defects that might exist within the immunoreaction spots on each xVFA sensing membrane. The immunoreaction spots functionalized with nine different capture antigens in duplicate are used as input for the deep-learning analysis. This helps to accurately analyze and interpret the results of the combined IgM and IgG antibody measurement for individual immunoreaction spots. The image processing algorithm can read and interpret two images per second, providing almost instantaneous results and ensuring a rapid outcome.

The decision neural network contains an input layer with M nodes (e.g., M = 25 with each immunoreaction spot on the multiplexed panel including positive and negative control spots), three fully connected hidden layers with 128, 64, and 32 nodes, in the first, second, and third layers, respectively. Each layer contains batch normalization, a 50% dropout, and a rectified linear unit (ReLU) activation function, except for the final output layer, which uses a sigmoid activation function, yielding a network output as a numerical value between 0 and 1. A final binary classification is then made by evaluating the numerical output with the blind cut-off value of 0.5. The training was carried out using a binary cross-entropy loss function with Adam optimizer along with a learning rate of 0.0001 and batch size 20. The architecture was determined by carrying out a grid search optimization of the major neural network hyperparameters including the number of layers, number of units per layer, regularization, dropout and learning rate.

Optimized neural network architecture was used during the selection of the optimal subset of immunoreaction spots. This spot selection process was implemented using a training set of serum samples (i.e., 40 serum samples from LDB) through the SFFS process, where the signals from each sensing channel were added one at a time into the input layer of the neural network and then trained via *k*-fold (*k* =5) cross-validation. After the addition of each input feature, the performance of the network was evaluated based on the area under the curve (AUC) scores and the input feature that yielded the best network performance for that iteration was then kept as an input feature until all 25 immunoreaction spots were included as an input (Figure 3d). The optimized model which utilized three peptides including modVlsE-FlaB, Var2FlaB and OppA4 was used for testing the remaining 10 serum samples from LDB and blinded validation of the assay platform using the CDC LD repository samples.

### Stability of peptides vs. proteins

To assess the stability of the antigen targets after immobilization on the nitrocellulose membrane, sensing membranes were spotted with 1 mg/mL of modVlsE-FlaB peptide and VlsE native *B. burgdorferi* protein and immobilized in three immunoreaction spots each. These membranes were then tested using control Lyme disease positive and healthy samples at various time points between days zero and ninety. The signal intensities corresponding to the peptides and proteins were compared to determine the stability of the antigens.

### Epitope mapping

Linear B cell epitope mapping was performed by ProImmune, Inc., using their ProArray Ultra Custom Microarray. Overlapping peptide libraires were generated for the indicated proteins consisting of 15-mer synthetic peptides overlapping by 10-amino acids (AA) (5AA-offset). The peptides were printed as arrays on glass slides, and the arrays were probed with 5 dilutions of serum from 8 patients considered highly seropositive for Lyme disease (9-10 bands positive on an IgG western blot). Positive binding was identified with fluorescently labelled anti-human IgM, IgG, IgA antibody (or just IgG for the paralogous proteins, BBA64, 65, 66, 73). An epitope was considered for further evaluation if a minimum of 6 of 8 sera (75%) demonstrated positive binding in two or more serum dilutions. Epitopes with high (>75%) sequence conservation among different strains of B. burgdorferi and low (<50%) sequence homology to unrelated antigens were prioritized. Prospective epitopes were synthesized as individual peptides of up to 45 AA in length (some epitopes overlapped multiple spots on the array). Epitope containing peptides were further screened by ELISA using larger serum sets containing serum from patients with EM lesions, Lyme arthritis, syphilis, rheumatoid arthritis, and serum from healthy volunteers living in Lyme disease endemic or nonendemic areas.

### Peptide selection using ELISA

The peptides were printed on glass slides and incubated with dilutions of 8 seropositive samples with early LD having at least 9-10 bands positive on the western blot. Epitopes with increased sequence conservation (>75%) among different strains of *B. burgdorferi* and low (<50%) sequence homology to unrelated antigens were prioritized. Serum antibody binding was visualized with fluorescently labeled anti-IgA, IgM, IgG, following which the slides were imaged and signal intensities were determined. Epitopes that bound antibodies in a minimum of two dilutions in at least 6 of 8 were synthesized as peptides. Epitope length was determined by the number of overlapping peptides identified in each analysis, but typically ranged from 15-35 AA.

### Peptide selection using xVFA

To screen the peptides selected using ELISA, we spotted 0.8 μL of 1 mg/mL of each peptide on a 25-spot sensing membrane in duplicates. We tested each peptide with all the samples from cohort 1 consisting of LD positive and healthy control samples. The signal intensities obtained were compared and the peptides were ranked based on their sensitivity, specificity, and correlation in identifying similar signatures of the patient samples.

### Clinical samples

Serum samples were obtained with unidentified labels with the sample information blinded until completion of the xVFA assay unless otherwise mentioned. The xVFA results were shared with the biobanks following which the labels were unblinded and identifiers were used to validate the performance.

Sample cohort 1 consisted of Lyme disease sera used for peptide screening by ELISA and were banked samples accumulated by Biopeptides, Corp. over the course of the last 30 years. They were collected from patients under informed consent with the approval of the institutional review boards of Stony Brook University in Stony Brook, NY, New York Medical College in Westchester, NY, and Gundersen-Lutheran Medical Center in La Crosse, WI. The 50 samples from Gundersen-Lutheran Medical Center in La Crosse, WI, had a clinician-documented EM lesion of > 4 cm, appropriate epidemiologic history (e.g., tick bite or exposure), and were seropositive according to a whole-cell ELISA (VIDAS by BioMerieux, Durham, NC, USA). We were not provided with the clinical laboratory results for the other deidentified Lyme disease patients beyond the fact that the patients were EM+. A total of 20 late Lyme disease samples were collected from patients at Gundersen-Lutheran Medical Center in La Crosse, WI, with Lyme arthritis (LA) (*n = 20*) that had one or more episodes of swollen joints, appropriate epidemiologic history, and positive reactivity using a whole-cell ELISA (VIDAS). The regions where the samples were collected, lower New York and Wisconsin, are highly endemic areas for Lyme disease. Sera from healthy volunteers were collected in New Mexico, which is not endemic for Lyme disease, and were purchased from Creative Testing Solutions (Tempe, AZ, USA). Sera from patients with rheumatoid arthritis (RA, rheumatoid factor status unknown) or who were Rapid Plasma Reagin positive (RPR+, first-tier test for syphilis) were purchased from Bioreclamation LLC (Westbury, NY, USA). These samples were collected in a region endemic to Lyme disease (the northeastern US). A total of 34 of the 35 RPR+ sera used in this study had positive or equivocal antibody levels against *Treponema pallidum* by ELISA (Abnova, Walnut, CA, USA). Some serum samples are not represented in all data sets because they were fully consumed during experimentation.

Sample cohort 2 included 12 clinical samples consisting of six Lyme disease positive samples from patients with different stages of LD purchased through a commercial vendor LGC diagnostics and six healthy control samples that were collected by the LDB from regions where LD is endemic.

Sample cohort 3 consisted of 50 samples from the LDB and was used for both training and testing of the deep-learning diagnostic model (Supplementary Table 3). The cohort included 25 laboratory-confirmed Lyme disease samples with signs and symptoms of early Lyme disease and were found to be positive by STTT. Additionally, 25 healthy control samples were collected from regions where Lyme disease is endemic. The LDB samples were characterized using screening ELISAs such as whole cell sonicates, C6 peptide ELISA, or VlsE/PepC10, and IgM and IgG western blots were run regardless of the first-tier results^29^. Samples were classified as either Lyme positive or negative using standard two-tier serology (STTT), and all 25 lab-confirmed samples were STTT positive while all endemic controls were STTT negative.

Sample cohort 4 included 32 samples acquired from the CDC’s Lyme serum repository research panel I, which consisted of 12 Lyme disease patient samples, 8 healthy control samples collected from regions where Lyme disease is endemic and non-endemic, and 12 look-alike disease samples which are known to be cross-reactive with LD diagnosis^30^. The CDC LDR samples were classified using both STTT and modified two-tier testing (MTTT). For STTT, the panels were measured using VIDAS Lyme IgM and IgG polyvalent assay by bioMérieux for first tier serology and MarDx Diagnostics, Inc IgM, and IgG immunoblotting assay for second tier serology. For the MTTT classification, tests from Zeus VlsE/pepC10, Zeus WCS EIA were used as first tier tests and Zeus IgM EIS and IgG EIA were used for second tier serology. No training was performed on the CDC dataset.

## Acknowledgements

The authors acknowledge Bay area Lyme Foundation’s Lyme disease biobank and the U.S. Centers for Disease Control & Prevention for graciously providing serum samples that were used for training and blinded validation of our platform respectively. We also acknowledge financial support from the National Institute of health (Grant #R44AI150060) and the National Science Foundation PATHS-UP Engineering Research Center (Grant #1648451).

