## Supplementary Information for "Single-tier point-of-care serodiagnosis of Lyme disease"

**Supplementary Information: Single-tier point-of-care serodiagnosis of Lyme disease**

**Contents**


- Supplementary Notes: Materials, preparation of antibody-AuNP conjugates, preparation of paper layers and ELISA procedure for paralogous proteins, BBA64, 65, 66, 73.

- Supplementary Table 1: List of secondary anti-human IgM and IgG antibodies screened for binding against anti-*Borrelia* antibodies.

- Supplementary Table 2: List of synthetic peptide antigens screened in this study.

- Supplementary Table 3: List of samples and their ground truth information as obtained from the Bay Area Lyme Foundation's Lyme disease biobank used in training of xVFA.

- Supplementary Table 4: Classification of samples obtained from the CDC's Lyme serum repository used for blinded validation of xVFA.

- Supplementary Table 5: Comparison of cost for xVFA test using recombinant protein antigens and synthetic peptides.

- Supplementary Figure S1: Screening of secondary antibodies for optimizing binding to anti-Borrelia antibodies.

- Supplementary Figure S2: Screening of BBA64-7, BBA65-94, BBA73 196-199 peptides using ELISA to measure interaction of synthesized peptides.

- Supplementary Figure S3: Sensitivity and specificity of individual peptides selected in the multiplexed panel without combination of analysis using a neural network.

- Supplementary Figure S4: Optimization of the xVFA to improve signal to noise ratio by tuning the buffer composition, IgM and IgG antibodies, sensing membrane porosity, AuNP size and sample volume.

**Supplementary Notes**

I. Materials

All Peptides listed in Supplementary Table 2 were synthesized by Lifetein (Hillsborough, NJ, USA). Dual epitope peptides were synthesized containing three glycine (G) residues between two epitope sequences. Recombinant OspC antigen was purchased from Prospec Inc. Anti-human IgM (#9022-01) and IgG (#9040-01) and anti-mouse IgG were purchased from SouthernBiotech (Birmingham, AL). Blocker™ Bovine Serum Albumin (BSA) (37525) was purchased from Sigma Aldrich. Nitrocellulose membranes (0.22μm (11327) and 0.45μm (11036)) were purchased from Sartorius Stedim North America Inc. A vivid plasma separation membrane (grade GX) was purchased from Pall Co., and the sample pad (CF7) as well as the conjugation pad (Grade Standard 14) were sourced from GE Healthcare Biosciences Corp. The absorbent pad (Whatman Grade 707) was acquired from OpticsPlanet, Inc. The gold colloidal solution (40 nm colloid, 15707-1) was purchased from Ted Pella, Inc. Foam tape (Super-Cushioning Food-Grade Polyethylene Foam Sheets 1/16") was purchased from McMaster-Carr.

II. Preparation of antibody–AuNPs conjugates

Complexes of mouse anti-human IgM/IgG on AuNPs were achieved by adding 100 μL 0.1M borate buffer (pH 8.4) and 20 μL of antibody (0.5 mg/mL) to 1 mL gold nanoparticle solution (40 nm, 1 OD) in a sterile Eppendorf tube. The mixture was incubated for one hour at room temperature, then 100 μL of 1% BSA in phosphate-buffered saline (PBS) was added as a blocking buffer. After blocking for 30 minutes at 25 ºC, the mixture was incubated at 4 °C for one hour. To remove excess mouse anti-human IgG/IgM, the conjugates were centrifuged at 4 °C for 15 min at 9000 rpm and washed 3 times by 1 mL washing buffer (10 mM Tris buffer (pH 7.2)). The supernatant was then re-suspended in 100 μL 0.1 M pH 8.5 borate buffer, containing 0.1% BSA and 1% sucrose. The final concentration of the antibody–AuNPs complexes was determined by optical density measurement at 525 nm using a well-plate reader (Synergy 2 Multi-Mode Microplate Reader, BioTek Instruments, Inc.). Finally the IgM/IgG-AuNP conjugates were diluted to 2 OD before mixing in 1:1 (IgM:IgG) ratio and used as the detection solution.

III. Preparation of functional paper layers and multiplexed sensing membranes

The paper layers were patterned using a wax printer (Phaser 8560, Xerox). The supporting layer and vertical flow diffuser were patterned on 0.22 μm NC membrane and 0.45 μm NC membrane respectively and incubated for 45 sec at 120 ºC in an oven to allow the printed wax to melt and diffuse downward into the nitrocellulose. The multiplexed sensing membrane was produced using a 0.22 μm NC membrane consisting of 25 spatially isolated immunoreaction spots defined by the wax printed barriers. After printing, the sensing membranes were incubated for 60 sec at 120 ºC in an oven to allow for the reflow of printed wax into the NC membrane. Each of the 25 immunoreaction spots were loaded with 1 mg/mL of the peptide antigen (18 spots) or 0.1 mg/mL of the Goat anti-mouse IgG solution (3 spots) and PBS buffer (4 spots). The spotted sensing membrane was allowed to dry in room-temperature for 30 minutes and then blocked with 1% BSA in PBS solution for 30 min to block non-specific binding, and again dried for 10 min at 37 ºC in a convection dry oven. The absorbent pad (1.2 × 1.2 cm), foam tape (1.7 × 1.7 cm for outside, 1.2 × 1.2 cm for inside dimensions), and asymmetric membrane (1.2 × 1.2 cm for absorption layer and 1st spreading layer and 1.4 × 1.4 cm for 2nd spreading layer) were all laser-cut (60W Speedy 100 CO2 laser from Trotec) to achieve precise dimensions.

IV. ELISA (for paralogous proteins, BBA64, 65, 66, 73)

96-well MAXISorp microtiter plates (thermofisher) were coated with 10μg/ml of peptide antigen in 0.1M Sodium Carbonate, pH9.4 for one hour at room temp. After one hour 1%BSA (Sigma) in PBS (blocking buffer) was added to each well and plates were incubated overnight at 4ºC. The following morning plates were washed 3 times with 0.05% Tween 20 in PBS using a Aquamax automated plate washer (Molecular devices), and 1:100 dilution of sera diluted in blocking buffer was added to each well. Plates were incubated at room temp for two hours and washed. To detect antibody binding, a 1:4000 dilution of AP-labelled goat-anti-human IgM or IgG (Southern Biotech) was added to each well and incubated for one hour at room temp. Plates were washed and developed using para-nitrophenyl phosphate (1mg/ml) in diethanolamine substrate buffer. Absorbance was quantitated at 405nm using a spectramax 384 plate reader (molecular devices).

**Supplementary Table 1.** Anti-human secondary IgM and IgG antibodies screened for binding against anti-*Borrelia* antibodies for the detection of LD infection.

| Code | Vendor | Host | Type | CAT# | Antibody |
|---|---|---|---|---|---|
| G1 | abcam | Mouse | Monoclonal | ab99770 | Mouse monoclonal [H2] Anti-Human IgG Fc |
| G2 | SouthernBiotech | Mouse | Monoclonal | 9040-01 | Mouse Anti-Human IgG Fc-UNLB (JDC-10) |
| G3 | | Mouse | Monoclonal | 9042-01 | Mouse Anti-Human IgG Fc-UNLB (H2) |
| G4 | | Goat | Polyclonal | 2014-01 | Goat Anti-Human IgG Fc, Multi-Species SP ads-UNLB |
| M1 | abcam | Mouse | Monoclonal | ab99741 | Mouse monoclonal UHB Anti-Human IgM mu chain |
| M2 | SouthernBiotech | Mouse | Monoclonal | 9020-01 | Mouse Anti-Human IgM-UNLB (SA-DA4) |
| M3 | | Mouse | Monoclonal | 9022-01 | Mouse Anti-Human IgM-UNLB (UHB) |
| M4 | | Goat | Polyclonal | 2023-01 | Goat Anti-Human IgM, Mouse/Bovine/Horse SP ads-UNLB |

**Supplementary Table 2.** List of all synthetic peptide antigens and their amino acid sequences used in this study.

| Synthetic peptide name | Amino acid sequence | Strain | Molecular weight |
|---|---|---|---|
| OspC | MTLFLFISCNNSGKDGNTSAC | B31 | 2223.53 |
| OppA | CYGQNWTSPENMVTSGPFKLKERIPNEKYVFEKNNK | B31 | 4277.88 |
| p35 | CDTGSERSIRYRRRVY | B31 | 2017.17 |
| ErpP | CKIEFSKFTVKIKNKD | B31 | 1928.33 |
| DbpA | CTILVNLLISCGLTGA | B31 | 1590.96 |
| DbpB | CKDLKNKILKIKKEATGKGVLFEAFTGLKTG | B31 | 3380.12 |
| BBK32 | CKKPMNKKGKGKIARKKGKSKVSRKEPYIHS | B31 | 3541.36 |
| BBA64 | CNESKKHKKEKRKGKV | B31 | 1927.31 |
| BBA65-94 | CTIKKISGGIRIQQGFEVA | B31 | 2048.45 |
| BBA65-101-02 | CKNTLEEIYNLIVDLTLIKKEW | B31 | 2679.19 |
| RecA164 | CIMFINQIRMRIGVMFGNPETTT | B31 | 2673.25 |
| RecA168-69 | CGGNALKFYSSLRLEVRKIEQVTR | B31 | 2768.25 |
| LA7 | CIPSKENAKLIVYFYDNVYAG | B31 | 2407.78 |
| FlilB | CVSRKGGLLPDIIIKI | B31 | 1725.18 |
| modClsE-FlaB | CVQEGVQQ EGAQQPGGGMKKNDQ IVAAIALRGVA | B31 | 3451.95 |
| Var2FlaB | CMKKTDNIAAAIVIRGVAKDGQFALKGGGVQEGVQQEGAQQP | B31 | 4312.97 |
| DbpA4-B6 | TILVNLLISCGLTGAGGGGGKDLKNKILKIKKEATGKGVLFEAFTGLKTGC | B31 | 5135.19 |
| BBA73-196-99 | MKRNKIWKTLKLFQITLLFSCSFYSKSNNTC | B31 | 3743.75 |

**Supplementary Table 3**. Samples obtained from the Lyme disease biobank and ground truth information.

| | *Clinical presentations* | | *Reference tests* | | | | *First-tier tests* | | | *Second-tier tests* | | *Diagnosis* | | |
|---|---|---|---|---|---|---|---|---|---|---|---|---|---|---|
| **Sample #** | ***EM/ Annular Rash*** | ***EM > 5 cm at Enrollment*** | ***B. burgdorferi PCR*** | ***B. miyamotoi PCR*** | ***Anaplasma PCR*** | ***B. Microti PCR*** | ***Whole Cell Lysate ELISA*** | ***C-6 Peptide ELISA*** | ***VlsE/PepC10 ELISA*** | ***WESTERN BLOT IgM**** | ***WESTERN BLOT IgG#*** | ***Seroconversion*** | ***Two-tier Positive*** | ***xVFA prediction*** |
| LDB01 | YES | NO | NEG | NEG | NEG | NEG | RE | POS | NA | POS | IT | NO | YES | POS |
| LDB02 | YES | YES | NEG | NEG | NEG | NEG | NR | POS | NA | POS | IT | NO | YES | POS |
| LDB03 | YES | YES | NEG | NEG | NEG | NEG | NR | POS | NA | POS | IT | NO | YES | POS |
| LDB04 | YES | NO | NEG | NEG | NEG | NEG | NR | POS | NA | POS | IT | NO | YES | POS |
| LDB05 | NO | NA | NEG | NEG | NEG | NEG | RE | POS | NA | POS | IT | YES | YES | POS |
| LDB06 | YES | YES | NEG | NEG | NEG | NEG | RE | POS | NA | POS | IT | NO | YES | POS |
| LDB07 | YES | YES | NEG | NEG | NEG | NEG | NR | POS | NA | POS | POS | NO | YES | POS |
| LDB08 | NO | NA | NEG | NEG | NEG | NEG | NR | POS | NA | POS | IT | NO | YES | POS |
| LDB09 | YES | YES | NEG | NEG | NEG | NEG | BL | POS | NA | POS | IT | NA | YES | POS |
| LDB10 | YES | NO | NEG | NEG | NEG | NEG | BL | POS | NA | POS | IT | NA | YES | POS |
| LDB11 | YES | YES | NEG | NEG | NEG | NEG | NA | POS | POS | POS | NEG | NA | YES | POS |
| LDB12 | NO | NA | NEG | NEG | NEG | NEG | NA | EQV | POS | POS | NEG | NA | YES | POS |
| LDB13 | NO | NA | NEG | NEG | NEG | NEG | NA | NEG | EQV | POS | NEG | NO | YES | POS |
| LDB14 | YES | YES | NEG | NEG | NEG | NEG | NA | POS | POS | POS | NEG | NA | YES | POS |
| LDB15 | YES | YES | NEG | NEG | NEG | NEG | NA | POS | POS | POS | NEG | NA | YES | POS |
| LDB16 | YES | YES | NEG | NEG | NEG | NEG | NA | POS | NA | NEG | POS | NA | YES | POS |
| LDB17 | YES | NO | NEG | NEG | NEG | NEG | NA | NEG | POS | POS | NEG | NA | YES | POS |
| LDB18 | YES | YES | NEG | NEG | NEG | NEG | NA | POS | POS | POS | POS | NO | YES | POS |
| LDB19 | NO | NA | NEG | NEG | NEG | NEG | NA | NEG | POS | POS | NEG | NO | YES | POS |
| LDB20 | NO | NA | NEG | NEG | NEG | NEG | NA | POS | POS | POS | POS | NO | YES | POS |
| LDB21 | YES | YES | NEG | NEG | NEG | NEG | NA | POS | POS | POS | NEG | NO | YES | POS |
| LDB22 | YES | YES | NEG | NEG | NEG | NEG | NA | POS | POS | POS | POS | NA | YES | POS |
| LDB23 | NO | NA | NEG | NEG | NEG | NEG | NA | POS | POS | POS | POS | NO | YES | POS |
| LDB24 | YES | YES | NEG | NEG | NEG | NEG | NA | POS | POS | POS | NEG | NO | YES | POS |
| LDB25 | YES | YES | NEG | NEG | NEG | NEG | NA | POS | NA | POS | NEG | NO | YES | POS |
| LDB26 | NA | NA | NEG | NEG | NEG | NEG | BL | NEG | NA | IT | IT | NA | NO | NEG |
| LDB27 | NA | NA | NEG | NEG | NEG | NEG | NR | NEG | NA | NEG | IT | NA | NO | NEG |
| LDB28 | NA | NA | NEG | NEG | NEG | NEG | NR | NEG | NA | NEG | NEG | NA | NO | NEG |
| LDB29 | NA | NA | NEG | NEG | NEG | NEG | NR | NEG | NA | NEG | NEG | NA | NO | NEG |
| LDB30 | NA | NA | NEG | NEG | NEG | NEG | NR | NEG | NA | IT | IT | NA | NO | NEG |
| LDB31 | NA | NA | NEG | NEG | NEG | NEG | NR | NEG | NA | IT | IT | NA | NO | NEG |
| LDB32 | NA | NA | NEG | NEG | NEG | NEG | NR | NEG | NA | IT | NEG | NA | NO | NEG |
| LDB33 | NA | NA | NEG | NEG | NEG | NEG | NR | NEG | NA | NR | IT | NA | NO | NEG |
| LDB34 | NA | NA | NEG | NEG | NEG | NEG | NR | NEG | NA | IT | IT | NA | NO | NEG |
| LDB35 | NA | NA | NEG | NEG | NEG | NEG | NR | NEG | NA | IT | IT | NA | NO | NEG |
| LDB36 | NA | NA | NEG | NEG | NEG | NEG | NA | NEG | NEG | NEG | NEG | NA | NO | NEG |
| LDB37 | NA | NA | NEG | NEG | NEG | NEG | NA | NEG | EQV | NEG | NEG | NA | NO | NEG |
| LDB38 | NA | NA | NEG | NEG | NEG | NEG | NA | NEG | NEG | NEG | NEG | NA | NO | NEG |
| LDB39 | NA | NA | NEG | NEG | NEG | NEG | NA | NEG | NEG | NEG | NEG | NA | NO | NEG |
| LDB40 | NA | NA | NEG | NEG | NEG | NEG | NA | NEG | NEG | NEG | NEG | NA | NO | NEG |
| LDB41 | NA | NA | NEG | NEG | NEG | NEG | NA | NEG | NA | NEG | NEG | NA | NO | NEG |
| LDB42 | NA | NA | NEG | NEG | NEG | NEG | NA | NEG | NA | NEG | NEG | NA | NO | NEG |
| LDB43 | NA | NA | NEG | NEG | NEG | NEG | NA | NEG | NEG | NEG | NEG | NA | NO | NEG |
| LDB44 | NA | NA | NEG | NEG | NEG | NEG | NA | NEG | NEG | NEG | NEG | NA | NO | NEG |
| LDB45 | NA | NA | NEG | NEG | NEG | NEG | NA | NEG | NEG | NEG | NEG | NA | NO | NEG |
| LDB46 | NA | NA | NEG | NEG | NEG | NEG | NA | NEG | NEG | NEG | NEG | NA | NO | NEG |
| LDB47 | NA | NA | NEG | NEG | NEG | NEG | NA | NEG | NEG | NEG | NEG | NA | NO | NEG |
| LDB48 | NA | NA | NEG | NEG | NEG | NEG | NA | NEG | NEG | NEG | NEG | NA | NO | NEG |
| LDB49 | NA | NA | NEG | NEG | NEG | NEG | NA | NEG | NEG | NEG | NEG | NA | NO | NEG |
| LDB50 | NA | NA | NEG | NEG | NEG | NEG | NA | NEG | NEG | NEG | NEG | NA | NO | NEG |

| | | | | |
|---|---|---|---|---|
| RE=Reactive | BL=Borderline | NR=Nonreactive | IT=Indeterminate | NA=Not applicable/Test not conducted |
| POS=Positive | NEG=Negative | YES=Positive clinical confirmation | No=Negative clinical confirmation | |

*Minimum of 2 of the 3 CDC specific bands (23,39,41 Kda) must be present. A western blot that does not meet the positive CDC criteria is considered "INDETERMINATE" by NYSDOH.

#Minimum of 5 of the 10 CDC specific bands (18,23,28,30,39,41,45,58,66,93 Kda) must be present. A western blot that does not meet the positive CDC criteria is considered "INDETERMINATE" by NYSDOH.

**Supplementary Table 4.** Samples obtained from the CDC and ground truth information.

| | *Standard two-tier tests (STTT)* | | | *Modified two-tier tests (MTTT)* | | | | *Diagnosis* | | | | |
|---|---|---|---|---|---|---|---|---|---|---|---|---|
| **Sample Group** | **VIDAS IgM/IgG EIA** | **IgM WB** | **IgG WB** | **Zeus VlsE/pepC10 EIA** | **Zeus WCS EIA** | **Zeus IgM EIA** | **Zeus IgG EIA** | **Standard Two-Tier prediction** | **MTTT IgM/IgG prediction** | **MTTT IgM prediction** | **MTTT IgG prediction** | **xVFA Prediction** |
| Early Lyme-EM Acute | 0/4 | 1/4 | 0/4 | 2/4 | 1/4 | 1/4 | 0/4 | 0/4 | 1/4 | 1/4 | 0/4 | 0/4 |
| Early Lyme-EM Convalescent | 4/4 | 4/4 | 1/4 | 4/4 | 4/4 | 4/4 | 1/4 | 4/4 | 4/4 | 4/4 | 1/4 | 4/4 |
| Late Lyme disease | 4/4 | 2/4 | 4/4 | 4/4 | 4/4 | 4/4 | 4/4 | 4/4 | 4/4 | 4/4 | 4/4 | 4/4 |
| Look-alike diseases | 0/12 | 0/12 | 0/12 | 1/12 | 1/12 | 1/12 | 0/12 | 0/12 | 0/12 | 0/12 | 0/12 | 0/12 |
| Healthy control | 2/8 | 0/8 | 0/8 | 0/8 | 3/8 | 3/8 | 0/8 | 0/8 | 0/8 | 0/8 | 0/8 | 0/8 |

**Supplementary Table 5.** Cost breakdown of xVFA test using recombinant proteins and peptide-based antigen panel.

| Types | Name | Cost/test ($) | |
|---|---|---|---|
| | | **Antigen assay** | **Peptide assay** |
| Paper materials | Asymmetric membrane | 0.38 | |
| | Interpad | 0.07 | |
| | NC membrane | 0.17 | |
| | Catridge (3D printed) | 1.00 | |
| Bio/chemical resources | Gold nanoparticles | 0.20 | |
| | Anti-Human IgG | 0.17 | |
| | Anti-Human IgM | 0.17 | |
| | Anti-Mouse IgG | 0.002 | |
| | BSA | 0.03 | |
| | Others | 0.40 | |
| | Antigens | **10.4** | |
| | Peptides | | **0.32** |
| | **Total cost per test** | **12.99** | **2.59** |

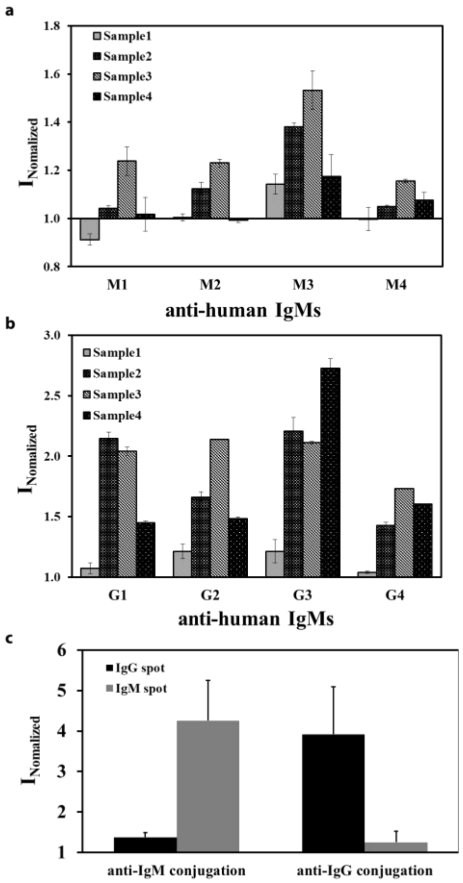


**Figure S1.** Screening of anti-human antibodies for binding to Lyme disease **a** IgM and **b** IgG antibodies. **c** Cross-reactivity evaluation of the IgM and IgG antibodies selected in the detection of a control LD patient sample containing anti-*Borrelia* IgM and IgG antibodies.

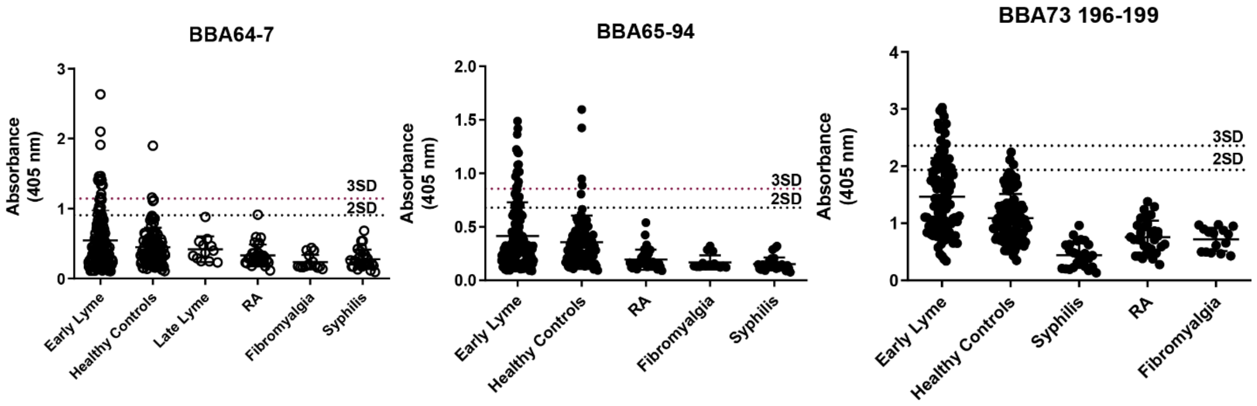


**Figure S2.** Screening of BBA64-7, BBA65-94, BBA73 196-199 peptides using ELISA to measure interaction of synthesized peptides with anti-*Borrelia* IgM antibodies tested against sample groups containing early Lyme disease patient samples, late Lyme disease, healthy control groups and cross-reactive diseases such as rheumatoid arthritis, fibromyalgia and syphilis. The cut-off for positivity was defined as three times the standard deviation of the healthy controls and an equivocal result was defined as a signal above two times the standard deviation. If the absorbance recorded for a healthy control sample was above three times the standard deviation, it was not considered for the cut-off determination.

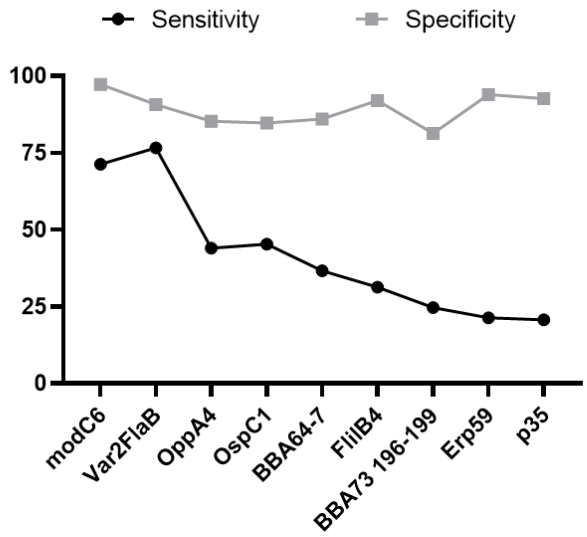


**Figure S3.** Sensitivity and specificity of individual synthetic peptides selected for diagnosing LD using xVFA without combining the signal intensities from reaction spots using a neural network model.

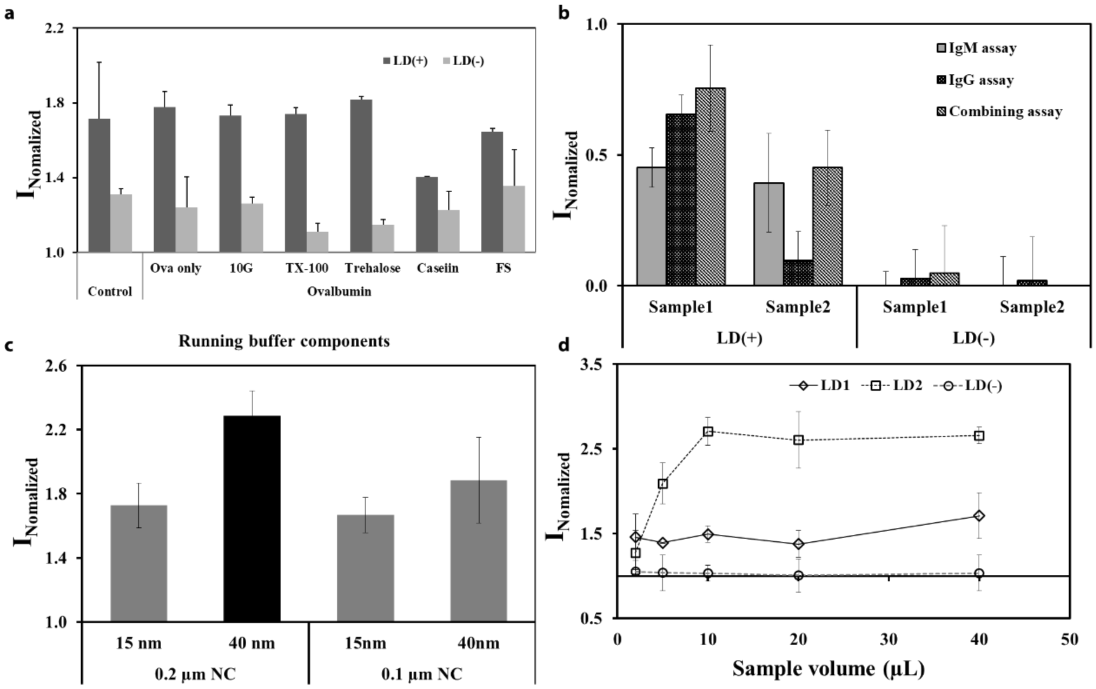


**Figure S4.** Optimization of the xVFA operation and assay reagents. **a** Evaluation of different running buffer condition where control represents a buffer containing 2% (w/v) ovalbumin, and 3% (w/v) Tween-20. Other compositions tested along with the control buffer were ovalbumin without the Tween-20 surfactant, 3% (w/v) 10G surfactant, 1% (w/v) Triton X-100, 2% (w/v) trehalose, 2% (w/v) casein and 10x dilution of Thermofisher Scientific superblock® buffer. Buffer containing 2% (w/v) ovalbumin, 3% (w/v) Tween-20 and 2% (w/v) trehalose was found optimal. **b** Testing of control Lyme disease patient samples with gold nanoparticles labelled with IgM only, IgG only and combined IgM and IgG secondary antibodies. **c** Screening of deposition of 15 nm and 40 nm gold nanoparticles on 0.2 µm and 0.1 µm nitrocellulose membrane. 40 nm gold nanoparticles deposited on 0.2 µm nitrocellulose membrane was found optimal. **d** Optimization of the sample volume added during the xVFA operation using two control Lyme disease patient samples and one control healthy sample. The signal was almost saturated at 10 µL sample volume and 20 µL was chosen as the volume of sample added for all clinical tests carried out.
